# qHTSWaterfall: 3-dimensional visualization software for quantitative high-throughput screening (qHTS) data

**DOI:** 10.1101/2022.06.15.496346

**Authors:** Bryan Queme, John C. Braisted, Patricia Dranchak, James Inglese

## Abstract

High throughput screening (HTS) is widely used in drug discovery and chemical biology to identify and characterize agents having pharmacologic properties often by evaluation of large chemical libraries. Standard HTS data can be simply plotted as an x-y graph usually represented as % activity of a compound tested at a single concentration vs compound ID, whereas quantitative HTS (qHTS) data incorporates a third axis represented by concentration. By virtue of the additional data points arising from the compound titration and the incorporation of logistic fit parameters that define the concentration-response curve, such as EC50 and Hill slope, qHTS data has been challenging to display on a single graph. Here we provide a flexible solution to the rapid plotting of complete qHTS data sets to produce a 3-axis plot we call qHTS Waterfall Plots. The software described here can be generally applied to any 3-axis dataset and is available as both an R package and an R shiny application.

## Introduction

Quantitative high-throughput screening (qHTS) was established over a decade ago as an approach to enable large-scale pharmacological analysis of chemical libraries.^1^ The method, an advance over the long-standing practice of testing compound libraries at a single concentration, was made possible by developments in assay technology, instrumentation, microtiter plate designs, and both commoditization and academic interest in chemical library generation.^2–5^ qHTS has been applied to enzymes, receptors, and biological processes using diverse libraries.^6–8^ For example, the National Center for Advancing Translational Sciences (NCATS), within the National Institutes of Health (NIH), has used qHTS in various aspects of drug and chemical probe discovery, including the evaluation of natural product extracts, drug repurposing, and drug combination testing.^9–11^

The large-scale acquisition of concentration-response curve (CRC) profiles has allowed a detailed study of the extent and structure-activity relationships (SAR) of chemotypes responsible for several confounding artifacts encountered in drug development. ^12–14^ The technique has also formed the basis of library toxicological profiling used in programs developing toxicity assessment methods.^15,16^ Furthermore, by exploring a chemical library spanning 4-5 orders of magnitude in concentration (e.g., nM to μM) relatively low potency starting points can be identified by including test concentrations far higher than previously considered.^6^

In addition to establishing a nascent library-wide SAR among the chemotypes in each library for the enzyme or phenotype under study, qHTS can provide insights related to a compound’s pharmacology. For example, in the work of Kinder et al., the CRC-derived Hill slopes from the qHTS of 4,500 drugs and investigational agents could be correlated with graded hyperbolic vs. ultrasensitive “switch-like” responses revealing a mechanistic basis for activity such as cooperativity or signal amplification (**Fig 1A-C**).^8^

**Figure 1.**
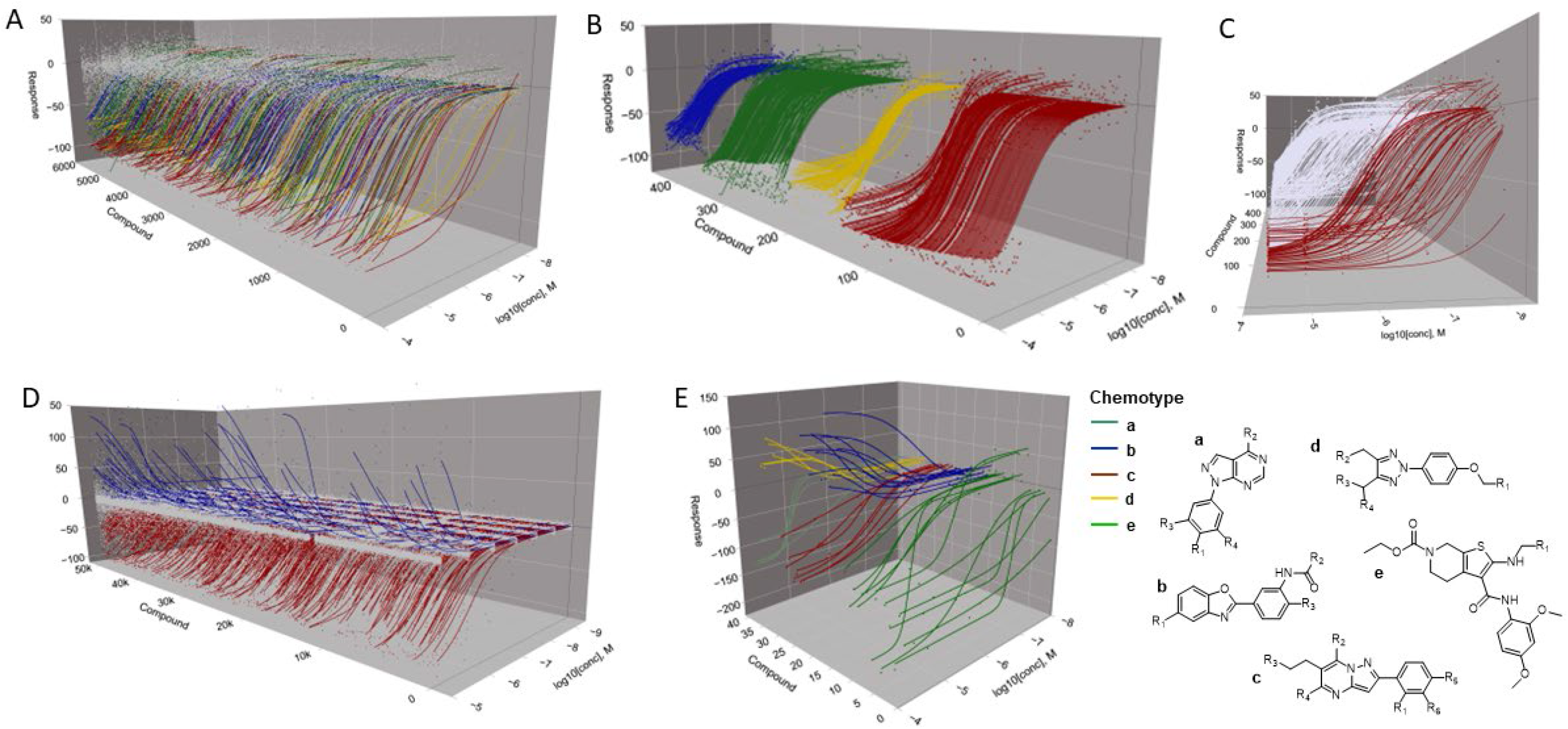
Various outputs for 3D visualization algorithm. **(A-C**) Graphs obtained using the one-output read algorithm. (**D**) Graph obtained using the two-output read algorithm. (**E**) Chemotypes a, c and e are associated with loss-of-signal response output, while chemotypes d and e display a gain-of-signal response as discussed in Martinez et al (ref 26). Data for graphs was obtained from the following PubChem AIDs, for plots in A-C: 1347405, 1347407 and 1347411; for plot D: 361; for plot E: 1508643.

Nevertheless, despite the increased use of this technique, delineating qHTS data remains challenging compared to the pairwise or two-axis graph types representing standard HTS data usually plotted as % activity vs compound ID.^17^ While large-scale and efficient twodimensional analysis for qHTS screening data has been developed, there remains a lack of 3-dimensional visualization tools for such libraries ^18–20^. In addition to providing a high-level overview of a qHTS experiment, three-dimensional graphs can allow the observation of patterns from thousands of CRCs not visible in two dimensions. For example, the output can be arranged and coded to highlight specific chemical and pharmacological properties embodied by the data, such as overall response efficacy (**Fig 1D**) as depicted in waterfall plot formats^21–24^ or related by structural chemotypes within the library (**Fig 1E**).

While the usage of qHTS has been increasing, few software packages can process the data to create three-dimensional graphs straightforwardly for chemical libraries on the order of 10s to 100s of thousands of members. With this in mind, we have developed an R package and associated application that creates three-dimensional graphs more efficiently than what is currently available in the market.

## Implementation

### Development

The 3D qHTS Waterfall Plot has been implemented in the R statistical programming language, using RStudio, and is developed as an R package to ease installation and use within developed R scripts and data analysis pipelines. The qHTSWaterfall package is also implemented as an R Shiny application so that in addition to R command line use, the application can be run through a user interface. The implementation can be installed on a user’s machine or hosted on a central Shiny Server instance as shown in **Figure 2**.

**Figure 2.**
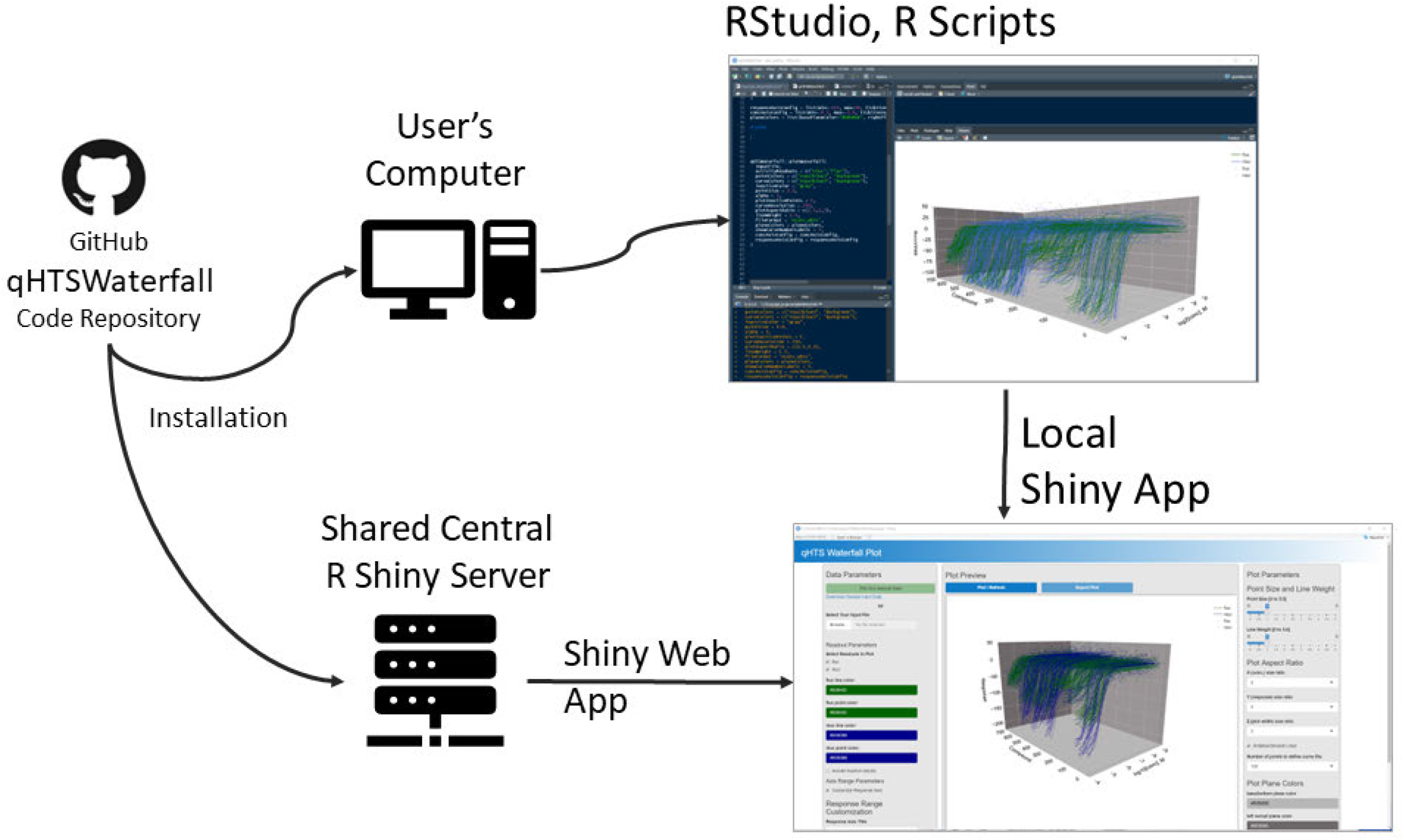
qHTSWaterfall Code Repository and Operating Environments.

## Results / Discussion

### Installation and Modes of Operation

The qHTSWaterfall package is implemented as an R package and as an R shiny application, having a user interface. Instructions for installation of the package can be found at our GitHub site in the readme section (https://github.com/ncats/qHTSWaterfall#readme) and are included in **Supplemental Figure 1**.

Starting the application using *runQHTSWaterfallApp()* in R will bring up a window with the application interface in the default application window. A button at the top of the interface allows users to move the application into an internet browser window, if desired. Clicking on the button labeled *Plot Our Sample Data* will access an included sample data set and plot the results. Note that the mouse scroller wheel or the zoom buttons in the upper right will allow one to zoom in and out. Other controls within the upper right context menu on the view, supported by the plotly package in R, allow one to pan and rotate the waterfall plot as well as capture the plot to a png image file.^26^ **Figure 3** shows the view of the qHTSWaterfall application user interface with the included sample data plotted, in this case having coincident reporter readouts of firefly luciferase (FLuc) and NanoLuc luciferase (NLuc).^25^ The plot controls are intuitive to use and include options to hide or show the various readouts, set colors for readout points and curve fit data, axis formatting, line weight, point sizes, and plot aspect ratio and background colors.

**Figure 3.**
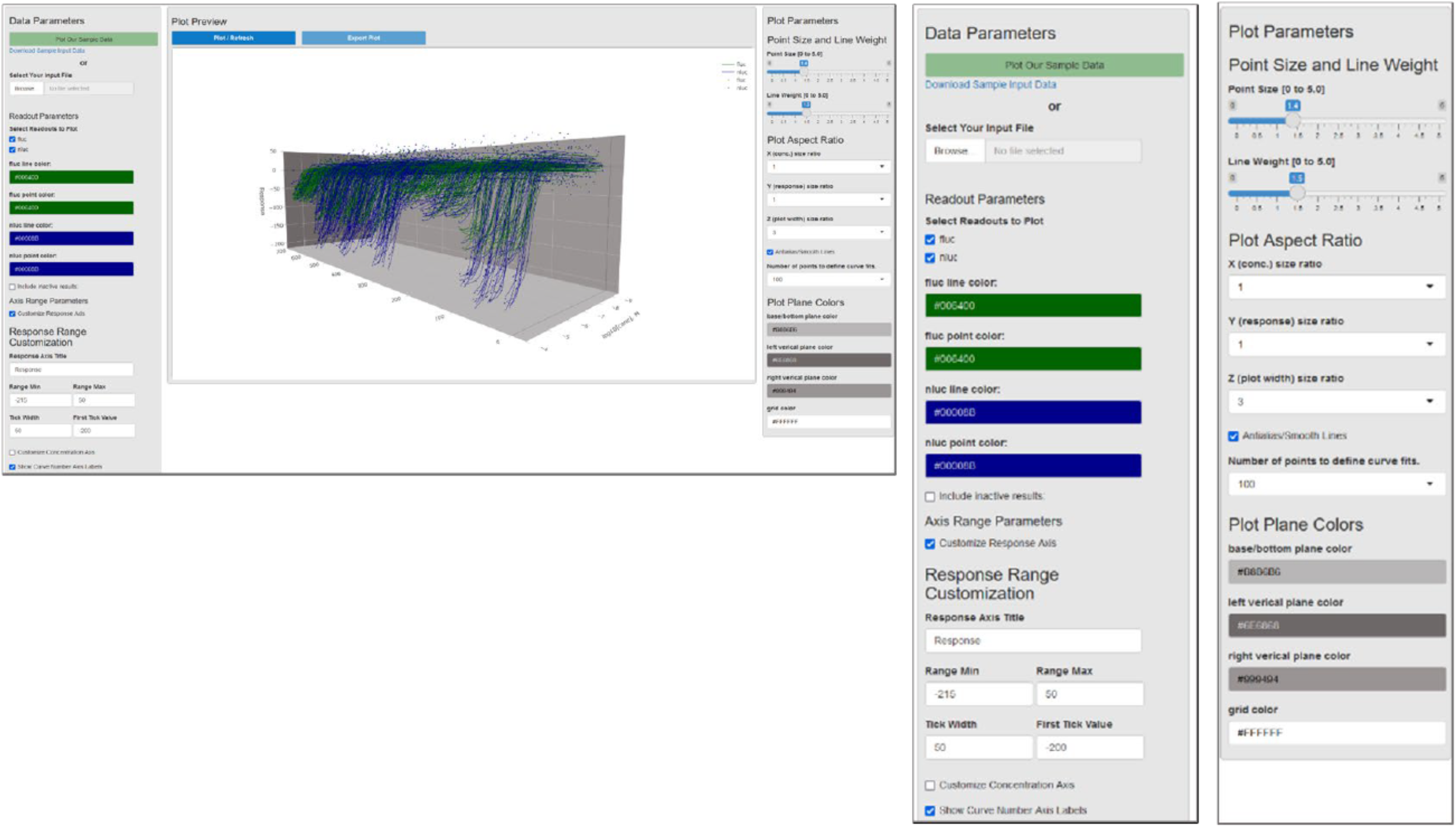
qHTSWaterfall interface showing a plot of sample data, hiding inactive results. The green and blue curves are individual coincidence reporter responses. Expanded on the right are the ‘Data Parameters’ and ‘Plot Parameters’ dashboards.

### Input File Format

Standard input file formats have been developed and sample files are available to plot and view within the application. These sample files can serve as templates for users’ input data. The software accepts comma-separated text files (.csv) or Microsoft Excel (.xlsx) files. The data within the files can be formatted in one of two forms. One format is specific to NCATS qHTS export format, however, most users will make use of the more generic format for general use, which is described in some detail here. A link in the upper left of the application will deliver an xlsx format sample input file with a color-coded header and notes on specific fields to include in the data. The header of this file, shown in **Figure 4**, illustrates the left (**A**) and right (**B**) columns of data, respectively.

**Figure 4.**
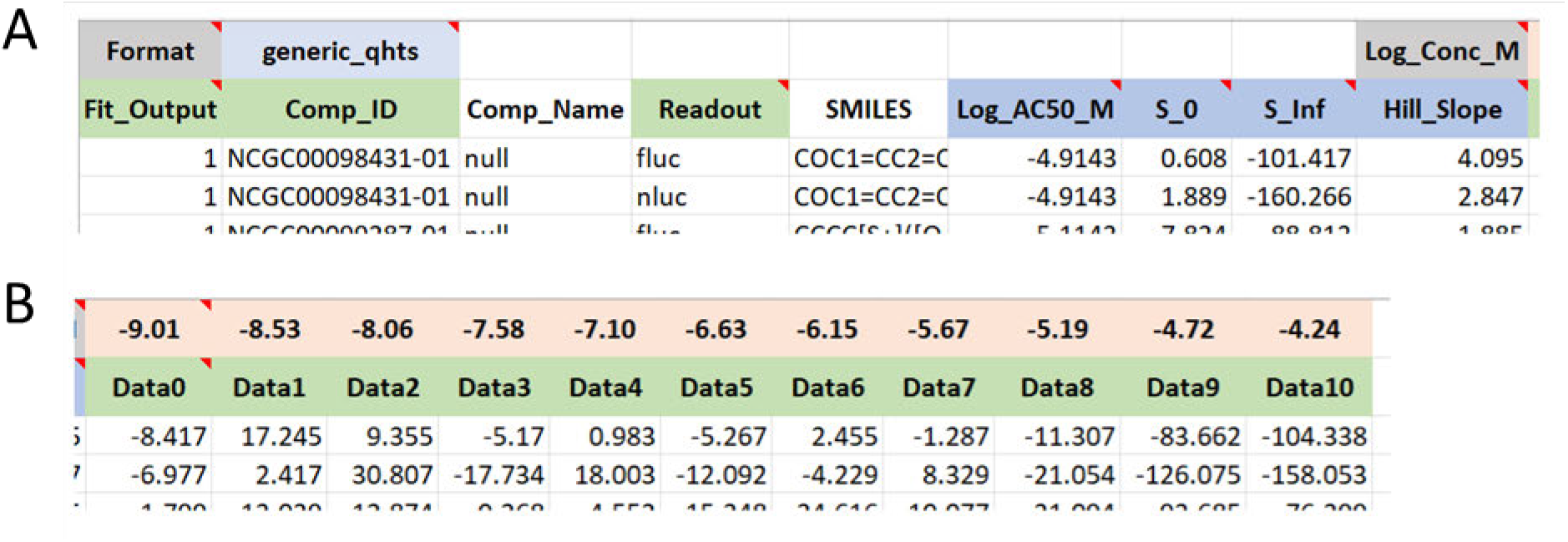
File format overview showing columns. (**A**). shows the format tag in the top row, compound annotations in columns 5, and concentration-response curve parameters in columns 6-9. (**B**). depicts example data columns, in this case an 11-point titration with log base 10 transformed molar (e.g., -pIC_50_) concentrations in the upper row, aligned with normalized data below.

The upper left of the file, referring to **Figure 4A**, has the keyword *Format* and the next cell to the right has the file format value. The value should typically be ‘generic_qhts’ unless working with NCATS format qHTS data in which this field will be ‘ncats_qhts’ and the format would be specific to NCATS qHTS format. The left-most column, Fit_Output, has values of 1 or 0, indicating if that compound response should be represented as a dose-response curve fit, or just by the data points that define that curve. Often users tend to only render full curves for active responses, those passing some level of curation, or responses that are of particular importance to show, such as results associated with a particular chemotype or readout type. Examples of this include a coincidence reporter response (**Figure 3**) where the two orthogonal reporter responses (FLuc or NLuc) are shown in green or blue, respectively.^26^ Another example would be a gain-of-signal vs loss-of-signal as shown in **Figure 1D**. Note that the order of compounds and associated response data will be preserved in the generated plot. This means that users can group compounds and responses based on activity criteria, readout type, chemical structure, or any other user-defined criteria (e.g., **Fig. 1A** vs **1B**). The *Comp_ID* column holds a user-supplied compound ID. Note that these IDs need not be unique, each compound can have multiple responses according to the specific *Readout* being reported on a particular row. The *Readout* column contains a descriptive name indicating the kind of response that is being reported on. In some assays, as shown, each compound may have different kinds of readout or even assay types. The readout column allows a compound to be represented more than once, to report on other measures of compound activity. In the sample file, a coincidence reporter assay reports on FLuc and NLuc outputs for each compound. Note here that within the application or R package, different *Readout* types can be shown or hidden, and point and line colors can be customized.

The curve fit parameter columns consist of those shown in light blue on the second row of **Figure 4A**, labeled *Log_AC50_M, S_0, S_Inf*, and *Hill_Slope*. These are the standard curve fit parameters associated with a four-parameter concentration-response curve fit against the Hill Equation. Please see **supplemental Figure 2** for the Hill Equation and explanation of the 4 associated parameters shown here.

The titration concentrations are captured in the input file, in the first row, just above the response values. Note that in **Figure 4A** upper right, we have a data tag, *Log_Conc_M*, to indicate the first column prior to the set of concentrations to read for data display. In **Figure 4B**, we show the primary data columns of the file. Each column has a specific log base 10 transformed molar concentration value and below that a data header labeled from Data0 to Data10, in this case. The input file can have any number of data columns but should use this naming convention to label starting at Data0.

Note that the 3D waterfall plot is constructed based on the order of compounds and their responses in the input file. This permits users to sort compounds based on a variety of criteria prior to plotting. Compound ordering can reflect structure-based clusters, response metrics such as potency and efficacy, readout type, or any combination of compound or compound response attributes. As an illustration of compound pre-sorting, **Figure 1B** features compound responses ordered and colored by NCATS curve class, a criteria-based response-curve classification system, and then ordered within each curve class by decreasing AC50.

Extra columns may be present in the file. In this example, we include a compound name and smiles structure string. Extra columns can be appended. The current restrictions are that the first two cells in the upper left should include the Format tag and the format value, and the data columns (Data0-Data_N_) and associated concentrations should be in a block of consecutive/contiguous columns as shown in **Figure 4B**.

## Conclusions

Obtaining a comprehensive view of bioactivity from a qHTS is highly informative from several perspectives. 3D data visualization can provide a high-level pharmacological assessment of overall library-assay activity allowing, for example, comparative analysis of assay activity vs library or *vice versa*.^1,7,8,10,12,13,26–28^ Further, by using specified sorting of compound similarity vs AC50, hill slope, max response, etc. highlighted information such as pharmacologic mechanism or chemical tractability can be conveyed to reveal actionable insights. For example, **Figure 1E** shows a plot from a qHTS follow-up where five firefly enzyme ligand chemotypes (*a-e*) are shown to have varying cellular consequences effects on firefly luciferase reporter output (PubChem AID=1508643).^26^

Producing overview plots for large screening campaigns had previously been a laborious process, using commercial software that were not optimally designed to handle this specific data and visualization type. The qHTSWaterfall application we present in this paper has allowed our lab to graph 3-dimensional qHTS data for various assays in a simple, and time-efficient manner. Generating overview presentations of qHTS data is roughly analogous to omics heatmaps in showing activity patterns over large data sets. To our knowledge, a free, opensource qHTS Waterfall plot software has not been previously available.

In addition, this program offers a facile means to generate a high-level analysis of the ever-increasing qHTS data appearing in repositories such as PubChem for anyone interested in studying a large and varied chemical biology data set. At the time this paper was written, there were over 15k HTS data sets in PubChem.^29^ While qHTS data can be represented by a 3-axis plot, the information content includes more than 3 parameters. For example, in addition to structural relationships among active compounds, each CRC contains pharmacologic parameters including an EC50 equivalent, a measure of potency, the hill slope, a mechanistic indicator, as well as the efficacy or magnitude of the response.

Our program allows biologists, chemists, informaticians, and the public to create 3-dimensional qHTS graphs clustered according to their preference as well as color aesthetics.

The user interface featured in the Shiny application helps users that are not proficient in R to produce plots, while others that wish to integrate the qHTSWaterfall plot into an existing R analysis workflow, can easily do so. 3-dimensional qHTS graphing allows researchers a general sense of trends, difficult to observe in a two-dimensional graphing format, relating to the interaction of chemical libraries with biological assays. Furthermore, this visualization can illustrate the scale of noise and artifacts between reporters and assays. In addition, our program allows scientists, regardless of previous programming experience, to create 3-dimensional qHTS data plots in an effective and timely manner. Researchers have the option to present data in clusters by the mechanism of action, activity, inhibition, or compound ID to organize data repositories such as PubChem.^29,30^

## Supporting information

Example Generic Format Data File

## Availability of Software and Data

Project Name: qHTS Waterfall (qHTSWaterfall R Package)

Project home page: https://github.com/ncats/qHTSWaterfall

Installation Instructions: https://github.com/ncats/qHTSWaterfall#readme

Operating systems: Platform independent

Programming language: R

License: Apache v2.0

Example data files: Included in the qHTSWaterfall R package or can be found in the source repository in this location:

https://github.com/ncats/qHTSWaterfall/tree/main/inst/extdata

https://github.com/ncats/qHTSWaterfall/raw/main/inst/extdata/Generic qHTS Format Example.xlsx

---

**Supplemental Figure 1.**
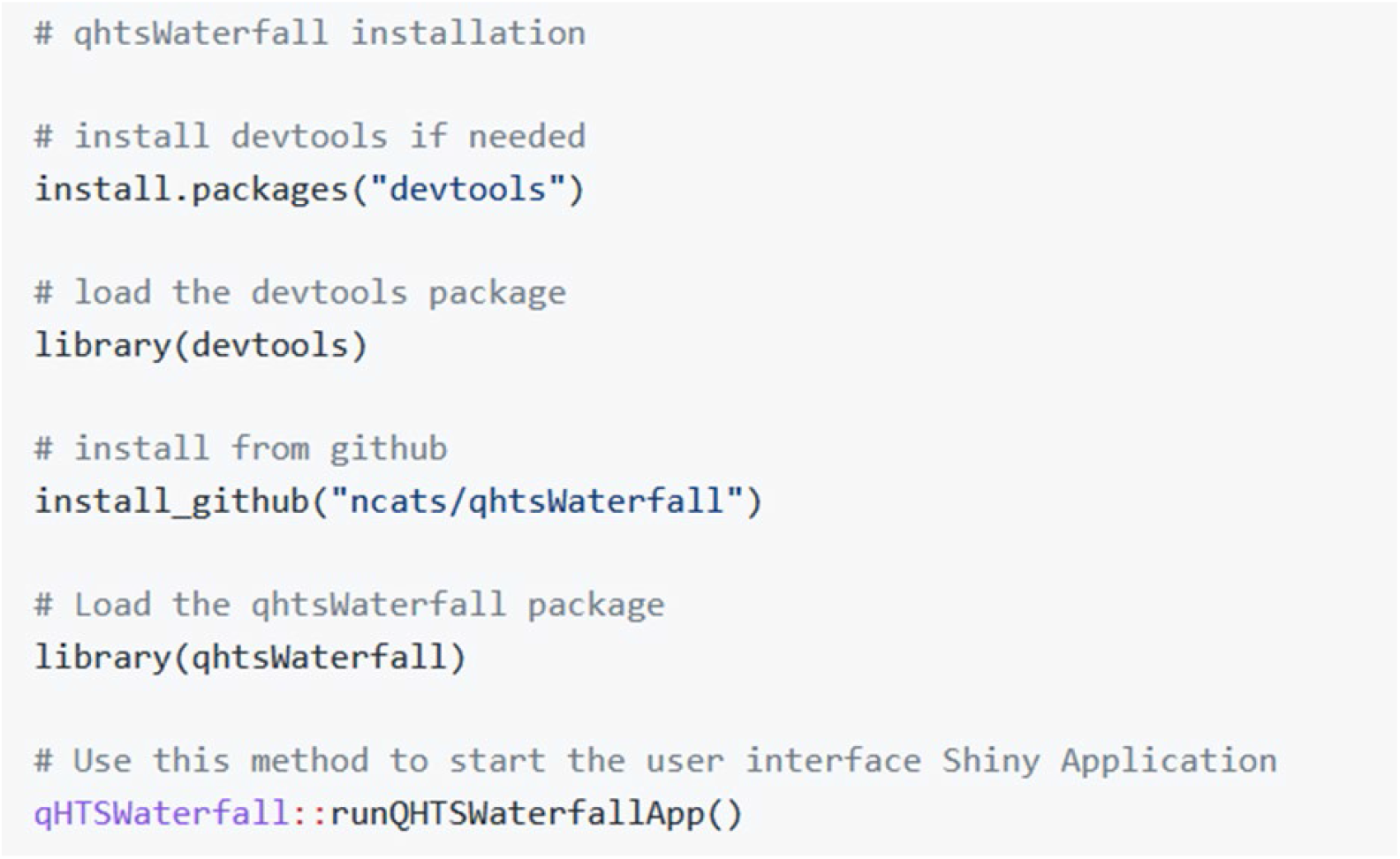
Instructions for installation and starting the qHTSWaterfall Application. The package *devtools* is required for installation from github.com and can be installed if needed.

**Supplemental Figure 2.**
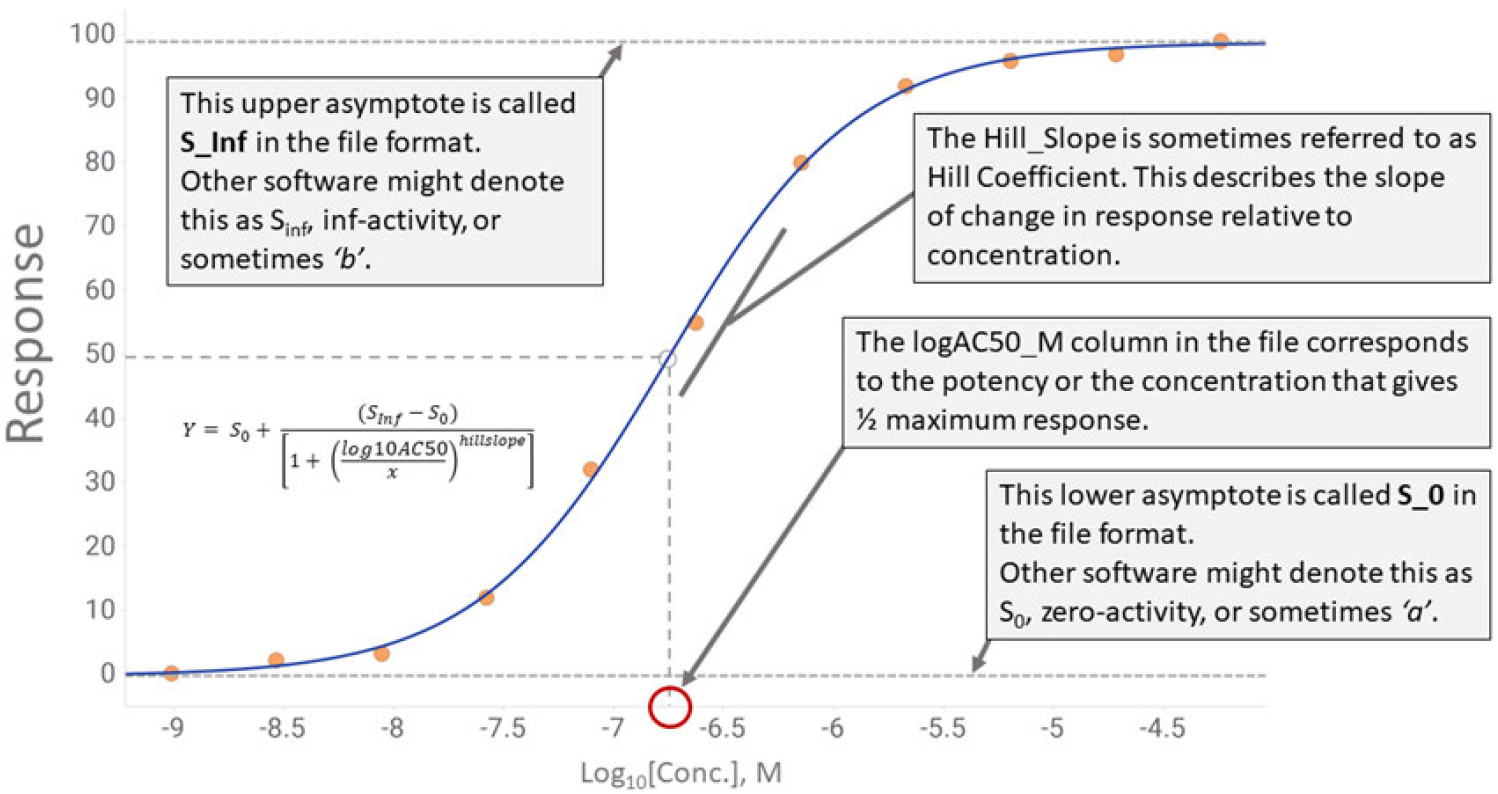
A sigmoidal concentration response curve. The 4 parameters contained in the input file (denoted in the file as S_0, S_Inf, Hill_Slope and logAC50) are explained here. Note that some software programs that generate these fit parameters may use different nomenclature to refer to these parameters.

